## Supporting Information for "Conserved Arginine Residues in Synaptotagmin 1 Regulate Fusion Pore Expansion Through Membrane Contact"

**Table S1**. Power Saturation parameters for spin labels in the arginine apex†

| Label Position | lipid | metal added | depth parameter (Φ) | position from lipid phosphate (Å) |
|---|---|---|---|---|
| C2AB 285R1 | Aqueous | none | –2.30 ± 0.009 | aqueous |
| | POPC:POPS | $Ca^{2+}$ | –1.62 ± 0.023 | –2.2 |
| | | EGTA | –1.76 ± 0.05 | –2.9 |
| | POPC:$PIP_2$ | $Ca^{2+}$ | –2.04 ± 0.02 | –4.7 |
| | | EGTA | –1.91 ± 0.03 | –3.5 |
| C2AB 349R1 | Aqueous | none | –2.41 ± 0.01 | aqueous |
| | POPC:POPS | $Ca^{2+}$ | –1.73 ± 0.075 | –2.0 |
| | | EGTA | –2.02 ± 0.064 | –4.5 |
| | POPC:$PIP_2$ | $Ca^{2+}$ | –2.14 ± 0.02 | –5.6 |
| | | EGTA | –2.22 ± 0.03 | –6.9 |
| C2AB 350R1 | Aqueous | none | –2.01 ± 0.03 | aqueous |
| | POPC:POPS | $Ca^{2+}$ | –1.76 ± 0.018 | –2.2 |
| | | EGTA | –1.83 ± 0.034 | –2.7 |
| | POPC:$PIP_2$ | $Ca^{2+}$ | –1.61 ± 0.02 | –1.1 |
| | | EGTA | –1.60 ± 0.05 | –1.1 |

† Depth parameters and approximate label positions obtained by progressive power saturation of the EPR spectrum (see text, Methods). Errors in the depth parameter are based upon standard deviations from at least 3 measurements.

**Figure S1**

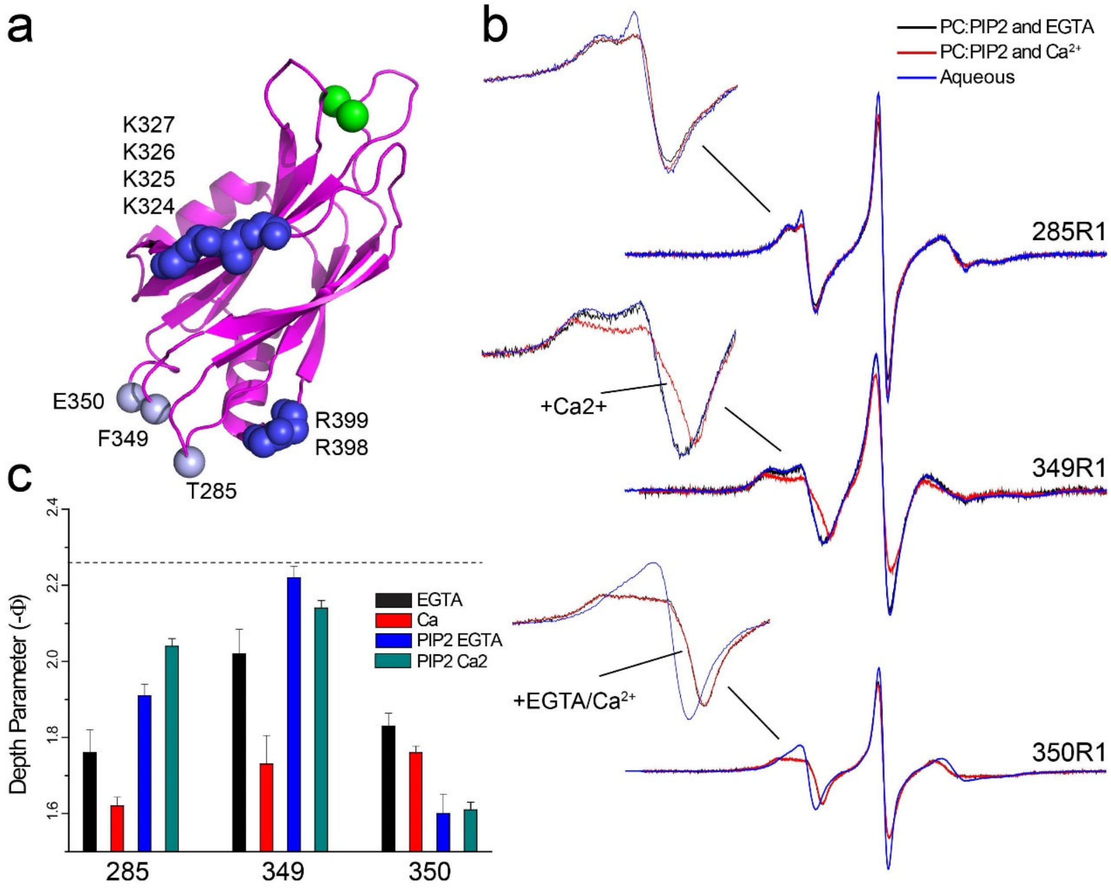


**Figure S1**. *The arginine apex of C2B contacts the membrane interface when membranes contain $PIP_2$.* **a**) Model for the C2B domain of Syt1 showing the labeled residues near the arginine apex (R399, R398) as well as lysine residues in the polybasic face (K324-327). **b**) EPR spectra from sites near the apex without membranes (aqueous, blue traces), and in the presence of PC:$PIP_2$ bilayers with $Ca^{2+}$ or EGTA (red and black traces, respectively). **c**) Membrane depth parameters comparing data for PC:PS with data for PC:$PIP_2$, with EGTA or $Ca^{2+}$. The effect of $Ca^{2+}$ is minimized in the presence of $PIP_2$, likely because the domain has a strong $Ca^{2+}$-independent association to $PIP_2$ containing membranes.

**Table S2**. Power Saturation parameters for spin labels in the arginine apex with RQ and RQRQ mutations†

| **Mutant/Label Position** | **lipid** | **metal added** | **depth parameter (Φ)** | **position from lipid phosphate (Å)** |
|---|---|---|---|---|
| C2AB 285R1 RQ | Aqueous | none | –2.28 ± 0.02 | aqueous |
| | POPC:POPS | $Ca^{2+}$ | –1.78 ± 0.028 | –2.3 |
| | | EGTA | –1.96 ± 0.05 | –2.9 |
| | POPC:$PIP_2$ | $Ca^{2+}$ | –2.13 ± 0.03 | –5.7 |
| | | EGTA | –2.04 ± 0.03 | –4.7 |
| C2AB 285R1 RQRQ | Aqueous | none | –2.26 ± 0.03 | aqueous |
| | POPC:POPS | $Ca^{2+}$ | –2.17 ± 0.008 | –6.2 |
| | | EGTA | –2.19 ± 0.02 | –6.4 |
| | POPC:$PIP_2$ | $Ca^{2+}$ | –2.23 ± 0.03 | aqueous |
| | | EGTA | –2.25 ± 0.05 | aqueous |
| C2AB 350R1 RQ | Aqueous | none | –2.08 ± 0.04 | aqueous |
| | POPC:POPS | $Ca^{2+}$ | –1.82 ± 0.02 | –2.6 |
| | | EGTA | –1.92 ± 0.03 | –3.5 |
| | POPC:$PIP_2$ | $Ca^{2+}$ | –1.64 ± 0.02 | –1.3 |
| | | EGTA | –1.66 ± 0.03 | –1.5 |
| C2AB 350R1 RQRQ | Aqueous | none | –2.07 ± 0.007 | aqueous |
| | POPC:POPS | $Ca^{2+}$ | –1.93 ± 0.02 | –3.6 |
| | | EGTA | –1.99 ± 0.03 | –4.3 |
| | POPC:$PIP_2$ | $Ca^{2+}$ | –1.66 ± 0.03 | –1.5 |
| | | EGTA | –1.68 ± 0.04 | –1.6 |

† Depth parameters and approximate label positions obtained by progressive power saturation of the EPR spectrum (see text, Methods). Errors in the depth parameter are based upon standard deviations from at least 3 measurements.

**Figure S2**

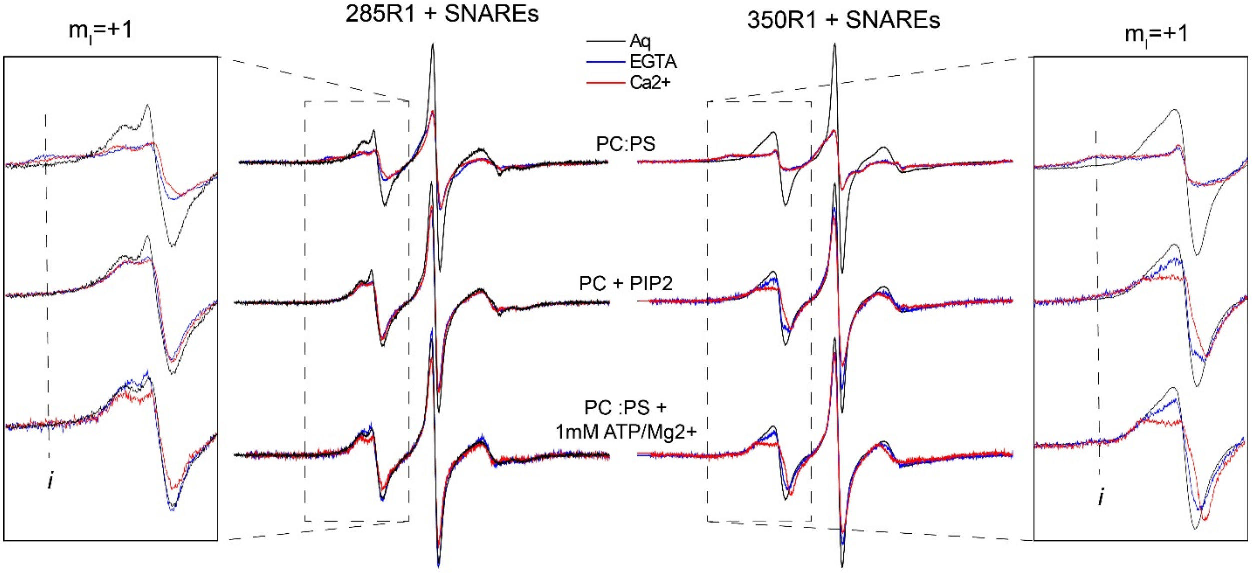


**Figure S2**. *EPR spectra from 285R1 and 350R1 indicate that either ATP or $PIP_2$ eliminate contact with membrane reconstituted SNAREs*. In PC:PS, contact with the SNAREs is evident from the appearance of an immobilized component in the region of the mI=-1 transition (see dashed line labeled "*i*") and the diminished normalized intensity of the EPR spectrum. However, when $PIP_2$ is present or when 1 mM ATP/$Mg^{2+}$ is present, the interaction is no longer observed.

**Table S3**. Power Saturation parameters for spin labels at the polybasic face of C2B†

| **Label Position** | **lipid conditions** | **metal added** | **depth parameter (Φ)** | **position from lipid phosphate (Å)** |
|---|---|---|---|---|
| C2AB 323R1 | Aqueous | none | –1.45 ± 0.05 | aqueous |
| | POPC:POPS | $Ca^{2+}$ | –0.804 ± 0.02 | 3.2 |
| | | EGTA | –1.38 ± 0.03 | 0.3 |
| | POPC:$PIP_2$ | $Ca^{2+}$ | –1.22 ± 0.02 | 1.21 |
| | | EGTA | –1.28 ± 0.03 | 0.89 |
| C2AB 329R1 | Aqueous | none | –1.66 ± 0.03 | aqueous |
| | POPC:POPS | $Ca^{2+}$ | –1.28 ± 0.03 | 0.88 |
| | | EGTA | –1.36 ± 0.02 | 0.47 |
| | POPC:$PIP_2$ | $Ca^{2+}$ | –1.19 ± 0.02 | 1.4 |
| | | EGTA | –0.298 ± 0.01 | 5.3 |

† Depth parameters and approximate label positions obtained by progressive power saturation of the EPR spectrum (see text, Methods). Errors in the depth parameter are based upon standard deviations from at least 3 measurements.

**Figure S3**

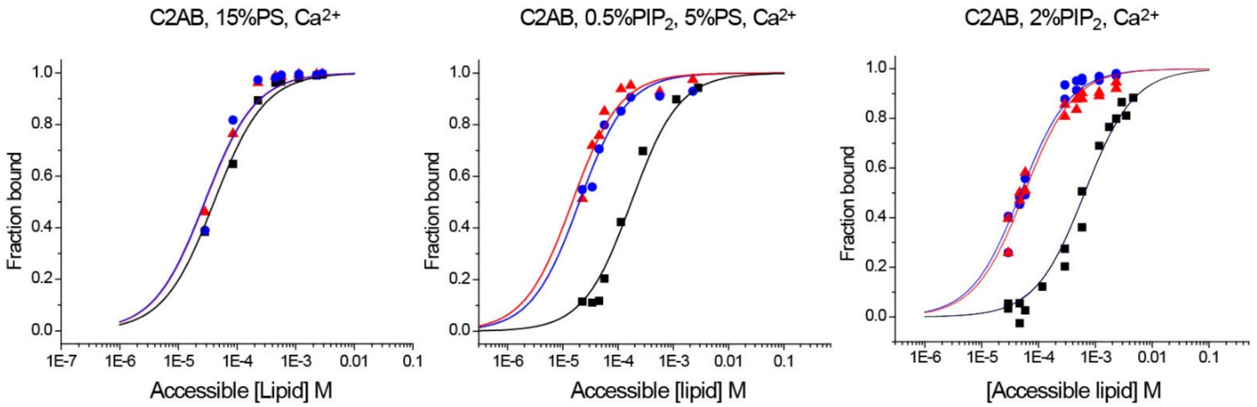


**Figure S3**. *Mutation of the polybasic face but not the arginine apex alters the membrane binding affinity of Syt1C2AB.* Equilibrium sedimentation data were obtained as described previously (1). Both the wild-type (blue circles) and RQRQ mutant (red triangle) have a similar binding affinity in the presence of $PIP_2$. Unlike the RQRQ mutant, the KAKA mutant in the polybasic face (black squares) exhibits a dramatically weakened membrane affinity.

**Table S4.** *SdFLIC results***.** The measured distances for each condition in $Ca^{2+}$ are reported from the mean of n experiments as $d_M \pm$ *standard errors*. Changes from this condition are reported as the mean of the changes from n experiments as *+/-(Δ* $d_M \pm$ *standard errors)*.

| C2AB added to | $d_M$ (nm)<br>+/- Δ $d_M$ (nm) | n |
|---|---|---|
| **Syx*192/SNAP-25/Syb1-96** in bPC/bPE/bPS/bPIP2/chol (34/30/15/1/20), in +100 μM $Ca^{2+}$ | 6.4±0.2 | 11 |
| +0.4 μM C2AB (WT) | + (4.7±0.2) | 6 |
| +0.4 μM C2AB (R398Q) | + (4.2±0.7) | 10 |
| +0.4 μM C2AB (R398Q/R399Q) | + (4.1±0.1) | 3 |
| +0.4 μM C2AB (K326A/K327A) | + (3.0±0.2) | 7 |
| **Syx*192/SNAP-25/Syb1-96** in bPC/bPE/bPS/chol (35/30/15/20), in +100 μM $Ca^{2+}$ | 6.1±0.4 | 25 |
| +0.4 μM C2AB (WT) | + (3.7±0.2) | 6 |
| **Syx*192/SNAP-25(AAA)/Syb1-96** in bPC/bPE/bPS/bPIP2/chol (34/30/15/1/20), in +100 μM $Ca^{2+}$ | 5.6±0.6 | 14 |
| +0.4 μM C2AB (WT) | + (2.1±0.5) | 7 |

**Table S5**. *Single SytKD-DCV fusion*. Percent fusion is calculated from the mean of n experiments. Errors are standard errors of repeats. Total number of docking and fusion events are the total numbers from all experiments.

| Acceptor SNARE membrane condition | Percent fusion | Total Number of Docking Events | Total Number of Fusion Events | n |
|---|---|---|---|---|
| Syx/SNAP-25 in bPC/bPE/bPS/bPIP2/chol (34/30/15/1/20), EDTA | 23±2 | 548 | 127 | 5 |
| +100 μM $Ca^{2+}$ | 24±2 | 2042 | 496 | 5 |
| +100 μM $Ca^{2+}$/0.4 μM C2ABwt | 50±5 | 1742 | 841 | 5 |
| +100 μM $Ca^{2+}$/0.4 μM C2AB (R398Q) | | | | |
| +100 μM $Ca^{2+}$/0.4 μM (R398Q/R399Q) | 45±2 | 440 | 199 | 5 |
| +100 μM $Ca^{2+}$/0.2 μM (K326A/K327A) | 35±2 | 431 | 153 | 5 |

**Table S6.** *Single WT-DCV fusion. Percent fusion is calculated from the mean of n experiments***.** Errors are standard errors of repeats. Total number of docking and fusion events are the total numbers from all experiments.

| Acceptor SNARE membrane condition | Percent fusion | Total Number of Docking Events | Total Number of Fusion Events | n |
|---|---|---|---|---|
| Syx/SNAP-25 in bPC/bPE/bPS/bPI/bPIP2/chol (25/25/15/4/1/30), | 41±1 | 823 | 340 | 14 |
| +100 μM $Ca^{2+}$ | 64±3 | 1753 | 1184 | 7 |
| Syx/SNAP-25 in bPC/bPE/bPS/bPI/chol (25/25/15/5/30), EDTA | 38±3 | 211 | 82 | 5 |
| +100 μM $Ca^{2+}$ | 47±3 | 386 | 183 | 6 |
| Syx/SNAP-25 in bPC/bPE/bPS/bPI/bPIP2/chol (32/32/15/1/20), EDTA | 26.6±2 | 516 | 134 | 5 |
| +100 μM $Ca^{2+}$ | 54.6±2.8 | 413 | 220 | 5 |
| Syx/SNAP-25(AAA) in bPC/bPE/bPS/bPI/bPIP2/chol (32/32/15/1/20), EDTA | 16.7±3.2 | 219 | 35 | 4 |
| +100 μM $Ca^{2+}$ | 33.6±3.3 | 185 | 63 | 4 |
